# A Platform for Oncogenomic Reporting and Interpretation

**DOI:** 10.1101/2021.04.13.439667

**Authors:** Caralyn Reisle, Laura Williamson, Erin Pleasance, Anna Davies, Brayden Pellegrini, Dustin W Bleile, Karen L Mungall, Eric Chuah, Martin R Jones, Yussanne Ma, Isaac Beckie, David Pham, Raphael Matiello Pletz, Amir Muhammadzadeh, Brandon M Pierce, Jacky Li, Ross Stevenson, Hansen Wong, Lance Bailey, Abbey Reisle, Matthew Douglas, Melika Bonakdar, Jessica M T Nelson, Cameron J Grisdale, Martin Krzywinski, Ana Fisic, Teresa Mitchell, Daniel J Renouf, Stephen Yip, Janessa Laskin, Marco A Marra, Steven J M Jones

**Affiliations:** Canada’s Michael Smith Genome Sciences Centre, Vancouver, BC, Canada; Bioinformatics Graduate Program, Faculty of Science, University of British Columbia, Vancouver, BC, Canada; QIAGEN Digital Insights, QIAGEN Inc., Redwood City, CA, USA.; Department of Medical Oncology, BC Cancer, Vancouver, British Columbia, Canada; Pancreas Centre BC, Vancouver, British Columbia, Canada; Department of Pathology and Laboratory Medicine, Faculty of Medicine, University of British Columbia, Vancouver, British Columbia, Canada; Department of Medical Genetics, University of British Columbia, Vancouver, BC, Canada; Department of Molecular Biology and Biochemistry, Simon Fraser University, Burnaby, BC, Canada

## Abstract

Manual interpretation of variants remains rate limiting in precision oncology. The increasing scale and complexity of molecular data generated from comprehensive sequencing of cancer samples requires advanced interpretative platforms as precision oncology expands beyond individual patients to entire populations. To address this unmet need, we created the Platform for Oncogenomic Reporting and Interpretation (PORI), comprising an analytic framework created to facilitate the interpretation and reporting of somatic variants in cancer. PORI is unique in its integration of reporting and graph knowledge base tools combined with support for manual curation at the reporting stage. PORI represents one of the first open-source platform alternatives to commercial reporting solutions suitable for comprehensive genomic data sets in precision oncology. We demonstrate the utility of PORI by matching 9,961 TCGA tumours to the graph knowledge base, revealing that 88.2% have at least one potentially targetable alteration, and making available reports describing select individual samples.

## Introduction

As the research and clinical applications of human cancer sequencing for precision medicine grow, there is an increased demand for the interpretation and reporting of genomic data in both research and clinical settings. Automation of cancer analysis research pipelines has improved the speed of reporting and the reproducibility of results. However, portions of the analysis remain refractory to automation. The human interpretation of genomic data remains one of the largest bottlenecks in comprehensive precision oncology ^1,2^.

To address this problem, a number of cancer knowledge bases have been created, including: OncoKB^3^; Clinical Interpretation of Variants in Cancer (CIViC)^4^; Cancer Genome Interpreter (CGI)^5^; Catalogue of Somatic Mutations in Cancer (COSMIC)^6^; MetaKB^7^; Jackson Laboratory Clinical Knowledge Base (JAX-CKB)^8^; Precision Medicine Knowledge Base (PMKB)^9^; My Cancer Genome^10^; Personalized Cancer Therapy (PCT)^11^; and Cancer Driver Log (CanDL)^12^. Despite the increasing availability of publicly accessible knowledge bases, these resources are distributed across a broad landscape of clinical and biological knowledge that is often disjointed and of varying structure. Integration of these tools into a reporting workflow to improve coverage^7^ is essential, yet left largely to individual users.

The increasing scale and complexity of the genomic and clinical data collected for sequenced tumour samples requires flexible analytic platforms suitable for automation^2^, both for the annotation of molecular profiles as well as the concise reporting of such information. While there are visualization tools ^13,14^ and commercial reporting applications available ^8,15,16^, there are few open-source reporting alternatives ^17,18^. Despite previous work demonstrating improvements in clinical comprehension of complex genomic data using interactive over static reports^19^, there are currently no open-source web applications for reporting in precision oncology. Open-source software is essential for promoting reproducibility and transparency in both research and healthcare, allowing the community to evaluate the softwares implementations and ensure their correctness ^20^. This is particularly important in research where the outcomes and insights will ultimately impact patient care. Furthermore, there is limited ability to build on and learn from closed source implementations within the research and clinical communities^21^.

While institutions have aimed to standardize workflows with respect to laboratory methods or even the bioinformatic tools used in variant calling^22^, reporting and annotation workflows remain diverse ^15^. By providing an open-source reporting platform that can be shared and improved by stakeholders, we aim to enable consistency in reporting and reduce redundancy in the development of individual bespoke tools. Here we present a novel research platform that integrates variant annotation through knowledge base matching into a precision oncology workflow and provides users a reporting interface to curate, edit, and interact with the resulting data. This improves knowledge translation and communication to the downstream medical professionals involved in the research. as well as facilitating scaling to include more patients through increased automation.

## Results

### Flexible Open Source Reporting with PORI

The Platform for Oncogenomic Reporting and Interpretation (PORI) was developed to facilitate the automated analysis of whole-genome and transcriptome sequencing data from human cancer samples to support precision oncology^23^ research initiatives (Figure 1a). The PORI platform consists of two main components: a knowledge base (GraphKB) and a reporting tool, Integrated Pipeline Reports (IPR).

**Figure 1.**
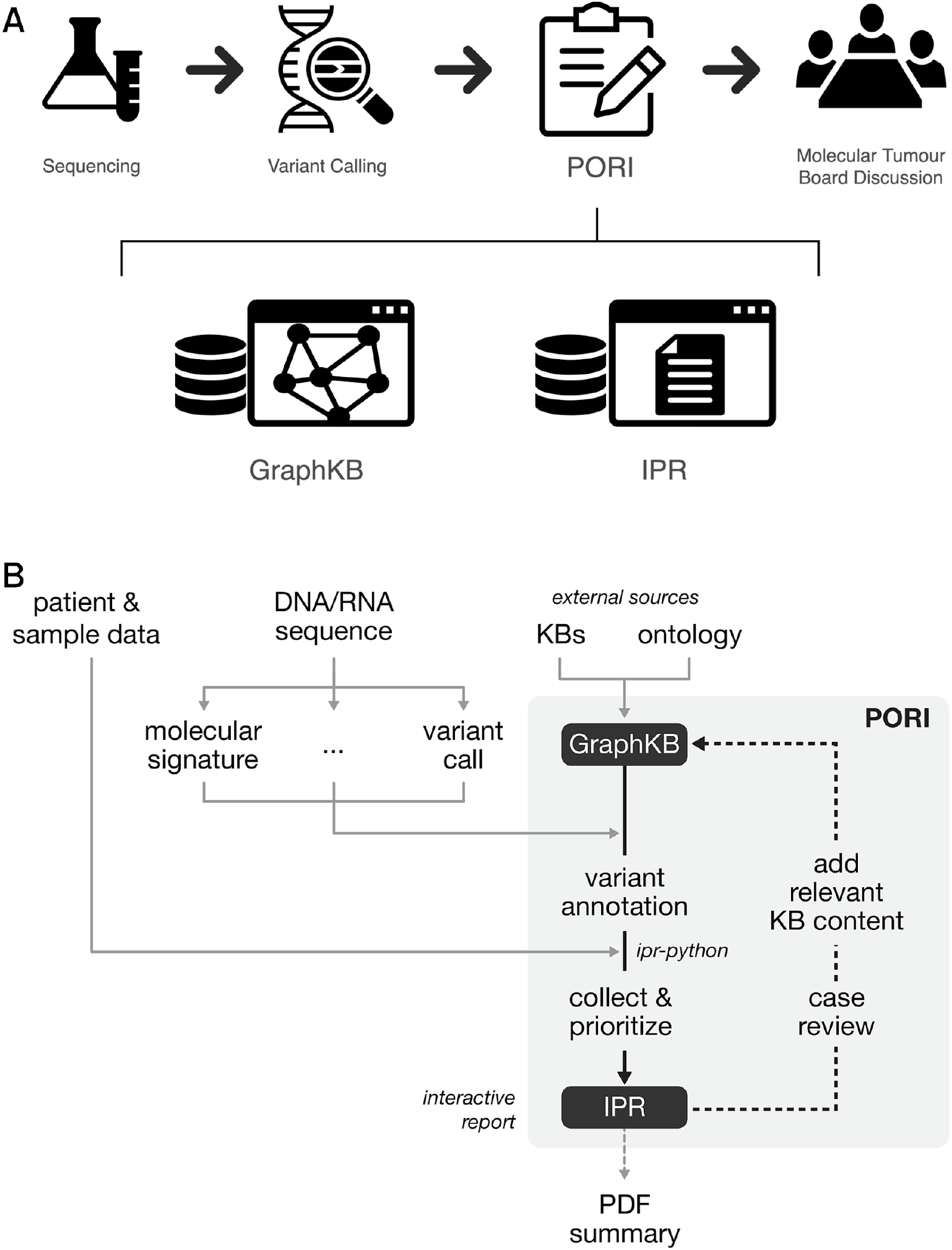
PORI Design showing both the placement of PORI within a precision oncology workflow (A) and the process of generating a report (B). PORI is used for the interpretation and reporting of genomic findings from tumour sequencing. Sequencing Data is taken as input to a number of bioinformatic pipelines and analyses. The results of these are loaded by the IPR report python adapter (ipr-python) and annotated with information from GraphKB. After annotation, the results are collated and prioritized based on matches for output into a report using the IPR interactive web platform. This is optionally manually reviewed by the case analyst who may add content to GraphKB as part of their literature review for the case and re-generate the report to include the newly added content. This report is shared with the molecular tumour board (MTB) to inform clinical decisions.

#### GraphKB: a graph database that incorporates diverse ontologies and domain knowledge

Traditional relational databases are ill-suited to storing hierarchical data or highly-related data like ontologies due to the prohibitively high cost of joining so many relations. In contrast, graph databases are designed with the connections between the data as a primary focus which allows complex relational queries to be performed efficiently^24^. This ability is leveraged heavily in GraphKB through the use of ontologies (Supplementary Figure 1). GraphKB is a graph database that has the ability to incorporate disease, drug, and gene ontologies and biological evidence statements from a large number of public external databases, including: HGNC^25^, Ensembl^26^, RefSeq^27^, Disease Ontology^28^, NCI Thesaurus (NCIt: https://ncithesaurus.nci.nih.gov), DrugBank^29^, Food and Drug Administration Substance Registration System (FDA: https://fdasis.nlm.nih.gov/srs), OncoTree (http://oncotree.mskcc.org), OncoKB^3^, CIViC^4^, CGI^5^, COSMIC^6^, ChEMBL^30^, Pubmed, and others. These ontologies are used as controlled vocabulary, but also to resolve redundant or related terminology through the linking of terms in and between ontologies. The GraphKB application stack includes two ways for the user to interact with the database, via the application programming interface (API) or the web client. Additionally, any queries made in the web client can display their API equivalent to help familiarize the user with the API (Supplementary Figure 2). GraphKB is primarily used to relate variants derived from patient data to known annotations in the literature. This is accomplished through the report python adaptor module included in PORI, ipr-python. The python adaptor collects and annotates the patient’s variants with information from GraphKB. This is then uploaded into IPR to create a report (Figure 1b; Supplementary Figure 3).

#### Integration of GraphKB and Integrated Pipeline Reports

The reporting component of PORI, IPR, is a web application for the visualization and dissemination of the genomic analysis and corresponding graphics, as well as evidence provided by the integration with GraphKB. It is used to review and communicate data both through the interactive web application as well as the production of portable document format (PDF) summaries (Figure 1b) suitable for dissemination of research reports to clinical personnel.

GraphKB and IPR are highly integrated. This integration is designed to facilitate the curation of clinically relevant content such as therapeutic biomarkers encountered during literature review of a patient’s variants. Reports are generated against a live version of GraphKB. Content relevant to a given case that was found through literature review and is not already curated in GraphKB can be added during case analysis and the report immediately re-generated. The quick turnaround time (~10 minutes) and minimal input requirements promote the updating of the knowledge base during case analysis which reduces the workload on the analyst by improving content coverage over time and consistency between reports. Additionally, inclusion of knowledge base entries into the report motivates the review of existing content; as the analyst reviews the report, they are linked from the report directly to the entries which have been matched in the knowledge base (Supplementary Figure 3). This ensures that relevant content is accurate and up to date, as it is reviewed and added with the highest priority.

In order to achieve a comprehensive understanding of a given patient's disease profile, the integration of diverse types of genomic alterations and complex signatures is required^31^. IPR collects output from many different types of bioinformatic analyses in a single report (Figure 1b). This provides the user with a central interface to interpret and interact with the data. To maintain flexibility, and recognizing the diversity of existing variant calling pipelines and workflows^22^, the PORI platform is run post-variant calling. In addition to the standard variant calls (SNVs, indels, structural variants, copy variants, RNA expression) PORI supports a number of other analyses including mutation signatures, tumour mutation burden, CIBERSORT^32^, MiXCR^33^ and OptiType^34^. A full list of the possible inputs to PORI can be found in the user documentation of the python report adaptor (https://bcgsc.github.io/pori_ipr_python).

Bioinformatics has a well-known software modality where tools are presented as proof of concept rather than production ready^35^. We have addressed this in PORI with standard techniques such as unit and integration tests using continuous integration and delivery systems. Additionally, PORI has been developed with multiple rounds of user testing. As a part of the Personalized OncoGenomics (POG) project, PORI has been refined based on feedback from three main user groups: clinicians, clinical trial nurses, and bioinformatic analysts ^36^. PORI has been used to generate and review 798 reports by 16 different authors covering 171 different diagnoses (Supplementary Figure 4).

### Improved Term Coverage through Multiple Ontologies

To avoid the issues associated with processing free text, most cancer knowledge bases choose ontologies as their source of controlled vocabulary (Supplementary Table 1). There are many competing ontologies and standards to choose from. The design of GraphKB follows previous work in integrating multiple external knowledge bases 7. However, GraphKB does not transform content upon input in order to harmonize between sources. GraphKB maintains the source form of the data and links entries using the underlying graph model instead. To accomplish this, GraphKB integrates multiple ontologies which are cross-referenced to one another using the links defined by each dataset (Figure 2).

**Figure 2.**
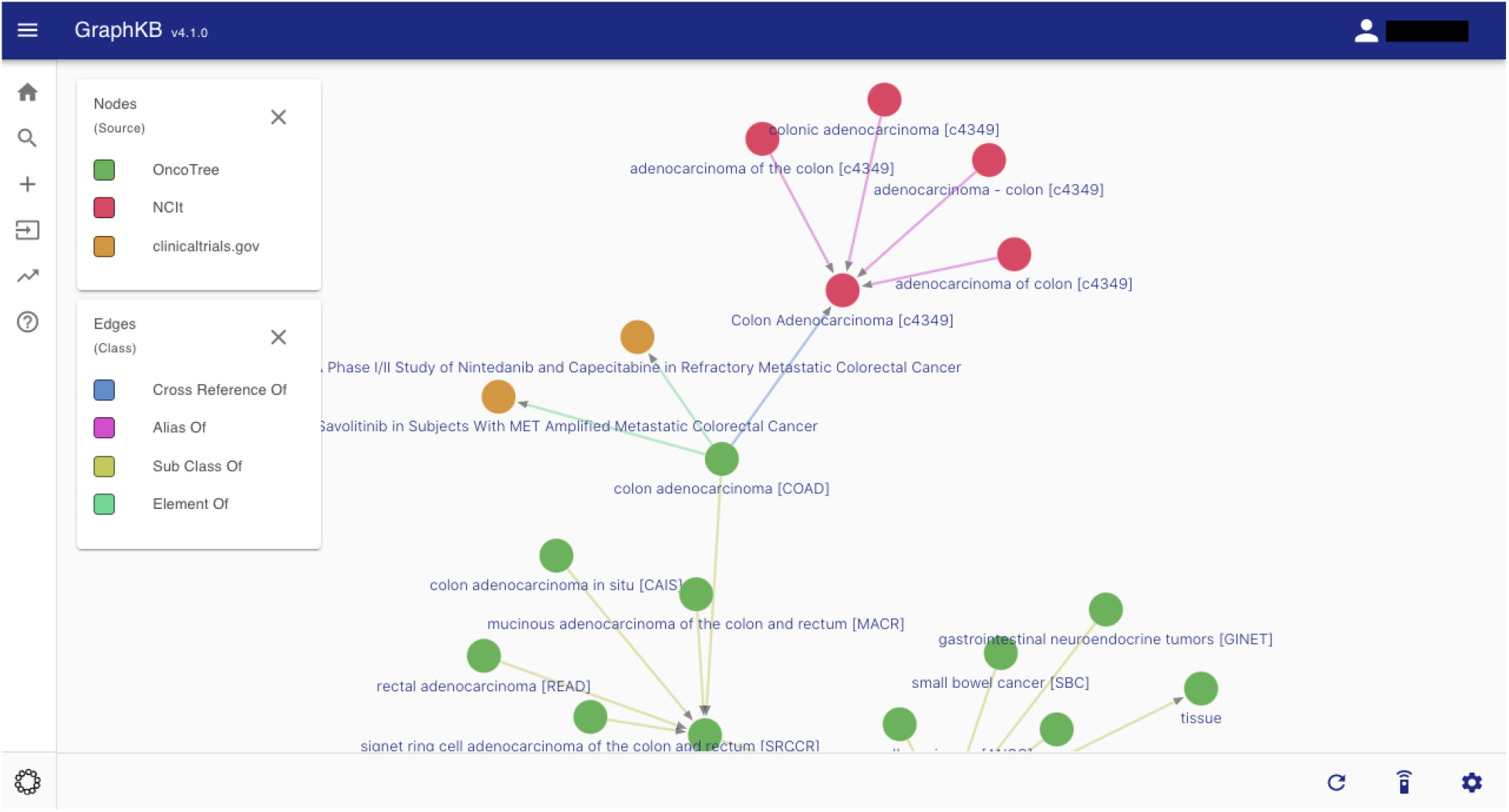
Graph View of content in the GraphKB web application showing a subset of links between different ontology terms for colorectal adenocarcinoma. Disease terms are shown from NCI thesaurus (NCIt) and OncoTree. For brevity, only a small amount of links to clinical trials (Clinicaltrials.gov) are shown.

To demonstrate the benefit of including multiple ontologies, widely used ontologies for diseases and drugs (Supplementary Table 1) were compared to determine overlap in coverage of terms as well as the total coverage of terms when ontologies were combined. Three disease definition resources were selected: OncoTree; Disease Ontology^28^; and NCIt. Four drug definition resources were selected: FDA SRS; DrugBank^29^; ChEMBL^30^; and NCIt. The full set of terms (indicated hereafter with a +) and the primary set of terms was calculated for each (Supplementary Table 2). The primary terms were considered to be the preferred terms and did not include aliases, synonyms, or product aliases but rather only terms which were given a unique identifier within the resource. By comparing common names between the full-term sets of each resource, we observed that more than 90% of disease and drug terms were unique to a single resource (Supplementary Figure 5). This indicates that the coverage of terms would be drastically reduced with the use of a single ontology.

The ability to leverage existing clinical resources represents one of the most enticing use cases of knowledge base content. Many of these resources do not use ontologies or even controlled vocabulary. In order to relate them to patient data, controlled vocabulary within the knowledge base is matched to the target vocabulary. To demonstrate an application of this, the terms from each ontology were compared to disease and drug terms listed in the ClinicalTrials.gov database (https://clinicaltrials.gov). The ClinicalTrials.gov database is a registry for clinical trials around the world that stores metadata regarding the trial such as location, eligibility criteria, and phase. Terms were extracted from the clinical trial records resulting in 116,237 therapy terms and 86,204 disease terms from 345,760 clinical trials. The terms from each resource were compared to the trial terms to determine the number of trial terms covered by each set of resource terms.

Terms used at a high frequency (100+) in ClinicalTrials.gov had higher coverage across ontology resources (Figure 3). The combined set of all terms, including synonyms, aliases and commercial product names, had the highest coverage (diseases: 0.63 and drugs: 0.96), with NCIt terms having the greatest coverage of any resource in isolation (diseases: 0.54; drugs: 0.95). However, when only primary terms were considered, ChEMBL (0.88) outperformed the other sources: FDA SRS (0.87); NCIt (0.72); and DrugBank (0.80). Similarly, when only primary terms were considered, the Disease Ontology (0.38) outperformed NCIt (0.14) and OncoTree (0.04). The 9% (7,758 diseases) improvement in disease term coverage of frequently used terms (100+) shows a clear benefit from the inclusion of multiple sources.

**Figure 3.**
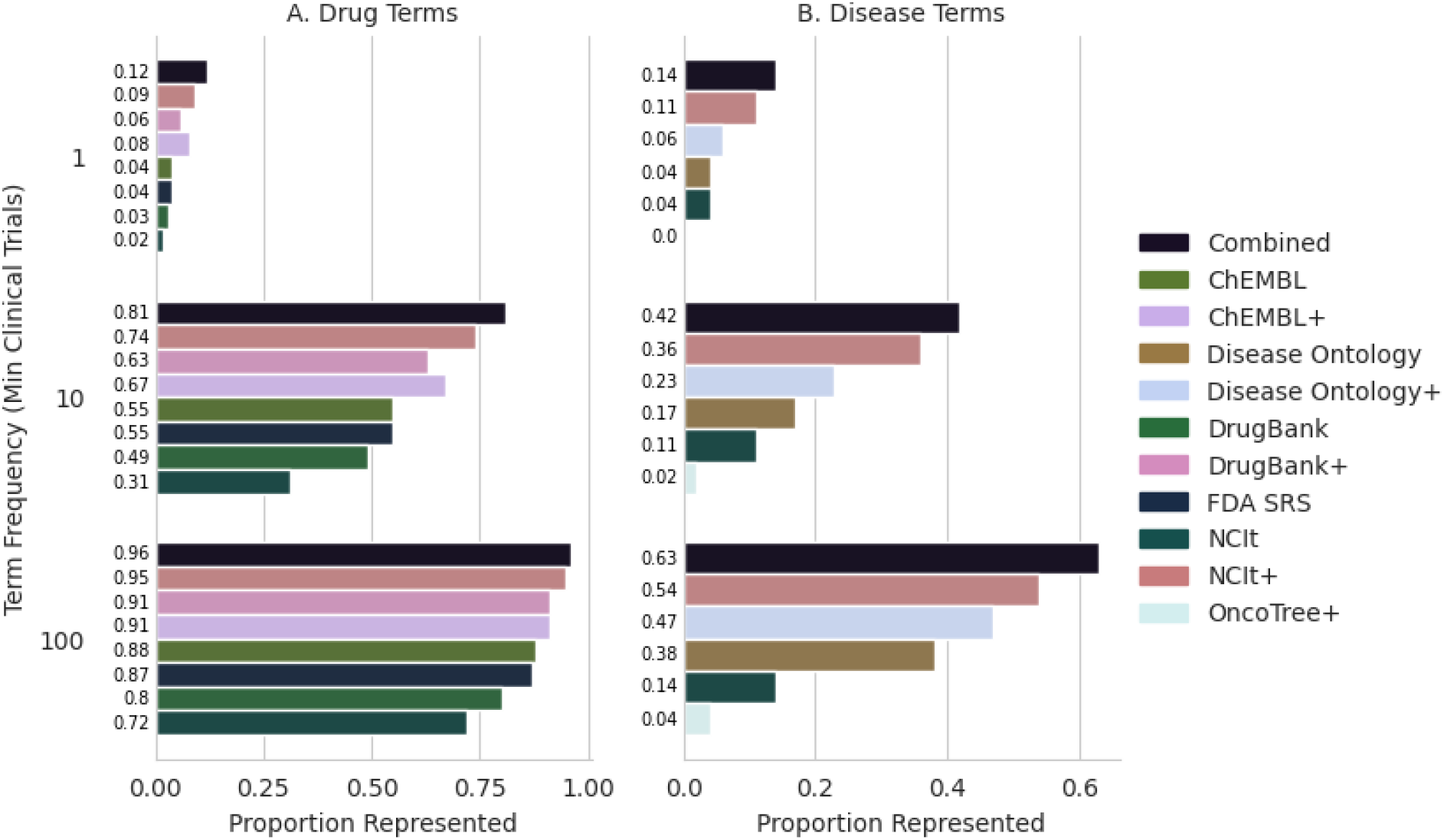
Coverage of Matched Drug (A) and Disease (B) Terms from Clinical Trials. A distinction was made between the primary/preferred terms for a given resource and the set of all terms (indicated with a +), which included synonyms, aliases, and commercial product names. Coverage was calculated for trial terms at 3 frequencies (1+, 10+, or 100+) where the frequency is calculated as the number of clinical trials a given term was used in.

### Leveraging the Graph Structure of GraphKB improves Concordance of Knowledge Base Sources

Due to the variability in the content and structure of cancer knowledge bases, it is necessary to integrate multiple cancer knowledge bases to ensure coverage of all relevant annotations^7^. GraphKB is able to support loading content from multiple external knowledge bases as well as adding content directly. Loading tools have been written for several popular knowledge bases which are included in the following knowledge base concordance analysis (Supplementary Table 3): OncoKB^3^; CIViC^4^; COSMIC^6^ (resistance mutations); Cancer Genome Interpreter^5^; and DoCM^37^. The content of these was compared to determine concordance between knowledge base sources. This was done at the level of conclusions, where conclusions are considered as the relevance (eg. sensitivity or resistance) and subject (ex. Drug or drug class) of a statement (Figure 4).

Before normalizing, there were 769 unique clinically informative (therapeutic, diagnostic, and prognostic) conclusions. After subject and relevance terms were normalized using ontology relationships, there were 696 unique conclusions, which demonstrates that while the different sources may appear initially to have disparate content, some of that content is in fact shared but done so with alternate representations such as aliases. By normalizing content using the graph model we are able to better quantify the levels of concordance.

**Figure 4.**
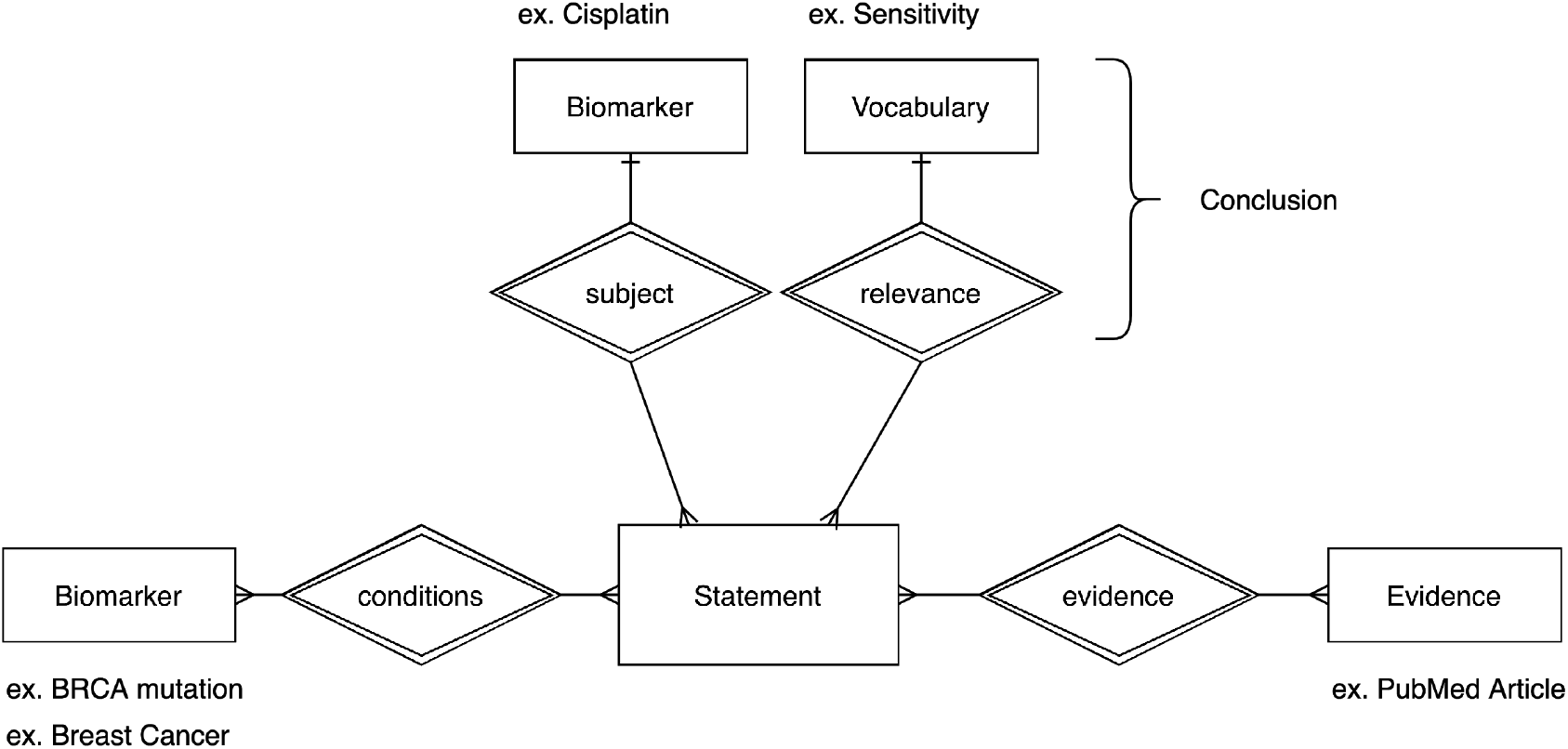
Statement Schema. Statements are composed of 4 main elements: conditions, subject, relevance, and evidence. A statement may be linked to any number of conditions but only one subject and relevance. The conclusion of a statement is considered to be composed only of the relevance and subject.

The agreement between knowledge base sources increased with normalization of related terms (Figure 5) from 14% (raw) to 19% (normalized) of conclusions shared in more than 1 source.

**Figure 5.**
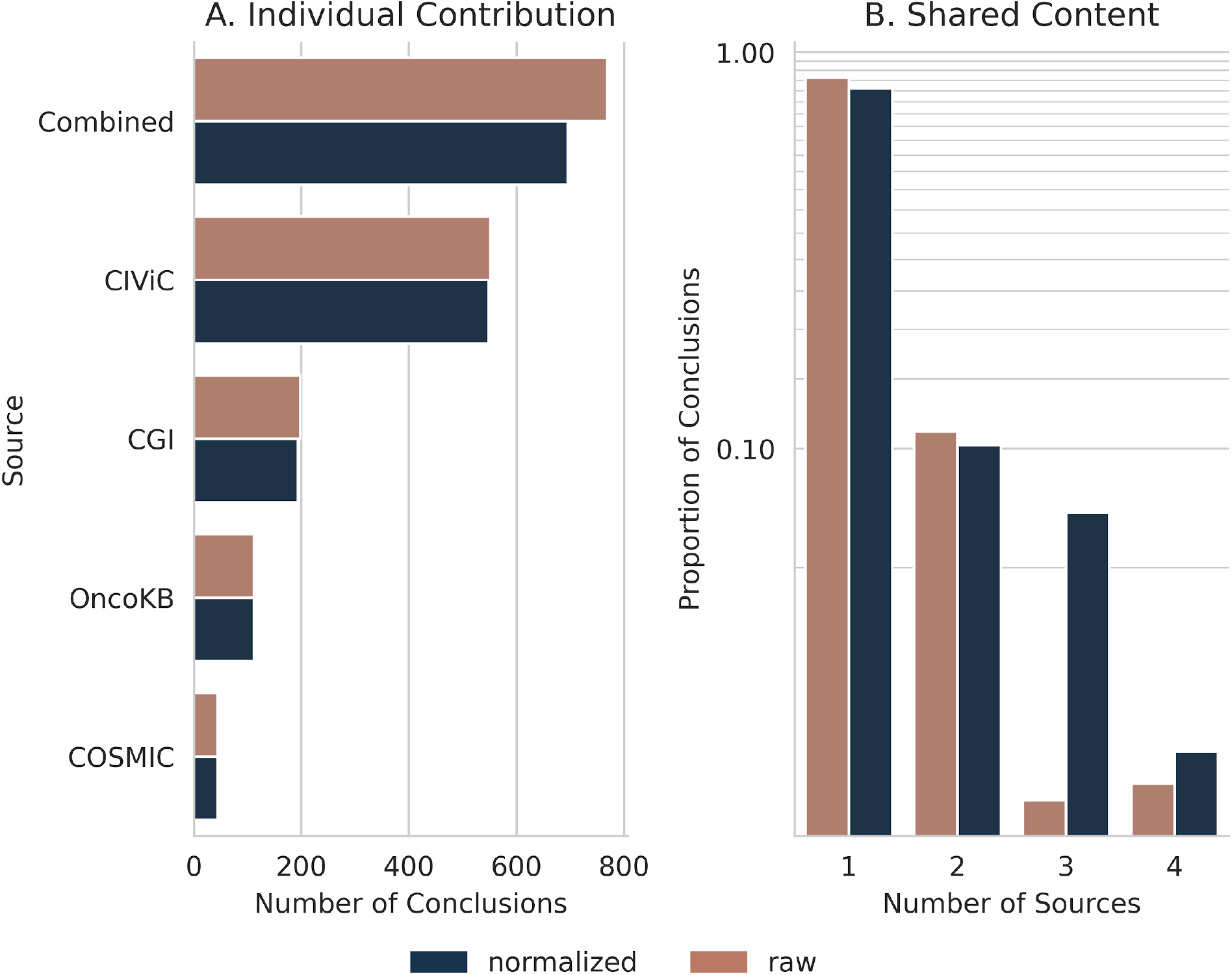
Clinically Informative Conclusion Agreement across knowledge bases. The individual contribution (A) of each source is shown as the number of unique conclusions which are given for both raw and normalized counts. The raw values represent the number of conclusions prior to normalization. The amount of content which is shared between sources (B) is shown as a fraction of the total number of unique conclusions.

### Application of PORI Using External Data Demonstrates the Benefit of Integration of Multiple Data Types

To demonstrate the flexibility of PORI both in using external data and supporting multiple data types, we analyzed the TCGA pan cancer atlas cohort^38^. Open-access data files were downloaded from cBioportal.org and analysed using the PORI platform ^39,40^. Mutations (mut); copy number variants (cnv); RNA expression variants (exp); and fusions (fus) from all studies were matched to GraphKB and annotated. Across all TCGA studies, there were 37,956 unique expression variants (20,136 increased expression and 17,820 reduced expression); 2,230,545 unique small mutations; 49,828 unique copy variants (24,906 amplifications and 24,813 deep deletions); and 21,291 unique fusions from 9,961 samples. There was a median of 21 unique conclusions per sample (1 sample per report), and a median of 13 conclusions related to therapeutic actionability (Supplementary Figure 6). Of these 9,961 samples, 88.2% (8,785) had variants which matched to one or more therapeutic conclusions. These cases were further analyzed as these represent potential therapeutic interventions or recommendations.

A large proportion of samples had therapeutic conclusions derived from a single variant type (small mutations: 18.1%; RNA expression: 11.4%; copy number: 5.4%), which demonstrates the importance of the inclusion of multiple variant types. If we only included a single variant type for GraphKB matching and reporting, then the number of samples where no therapeutic conclusions were found would increase by a minimum of 25.1% (2,495 samples) depending on which variant type was selected (Figure 6).

**Figure 6.**
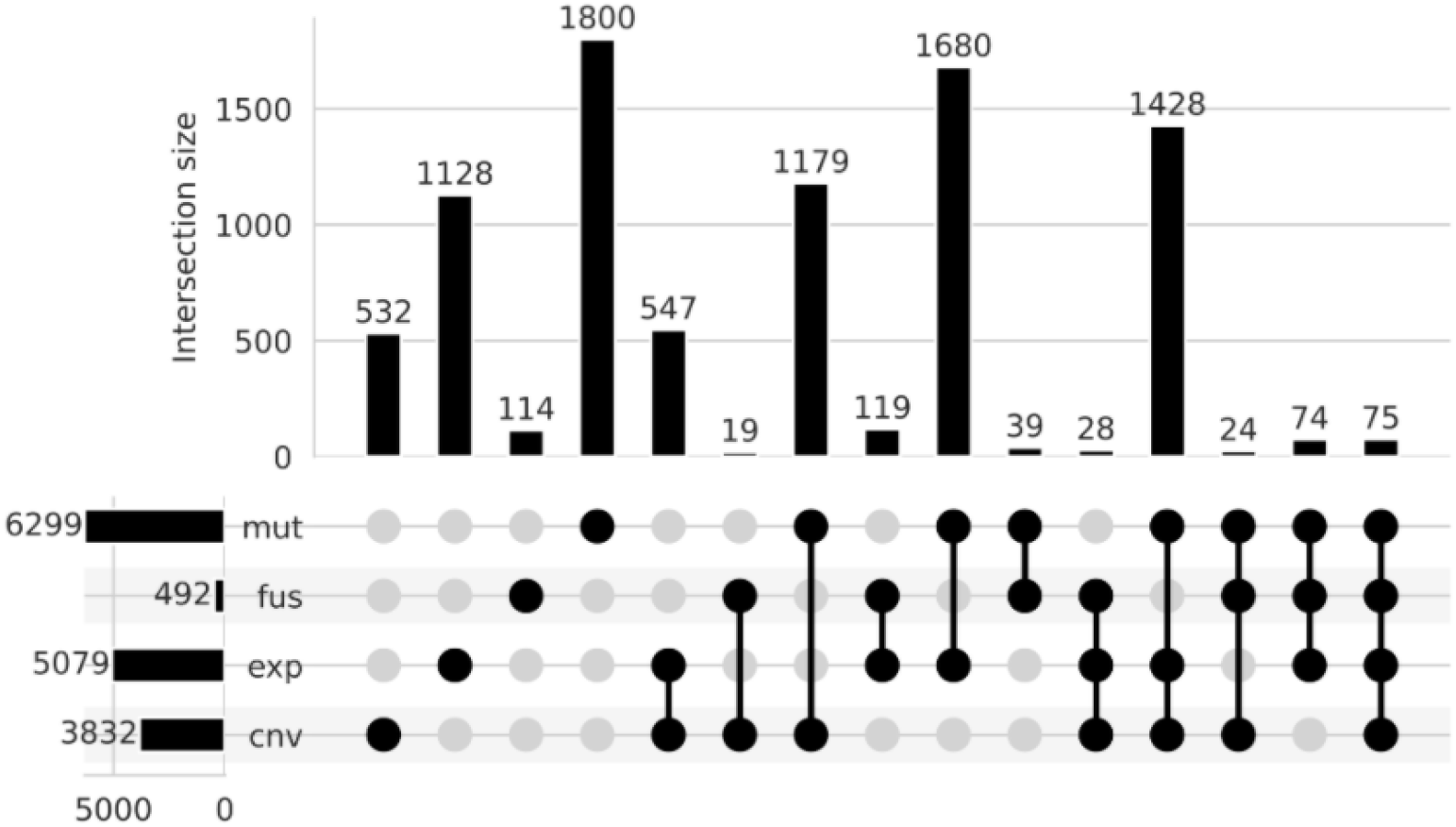
Proportion of samples with therapeutic conclusions derived from each combination of variant types. Upset plot of the number of samples with therapeutic conclusions from annotation of a given variant type. Sample variants are divided into four types: copy variants (cnv); single nucleotide variants and indels (mut); gene fusions (fus); and RNA expression (exp) variants. The left-hand bar plots are the total number of samples which have 1 or more therapeutic conclusions matched to the listed variant type. The upper bar plots show the number of samples in each of the intersection groups. These groups are mutually exclusive.

As expected, since small mutations have the greatest coverage across all public knowledge base sources used in this analysis, we observed the greatest contribution to therapeutic conclusions from statements associated with this variant type (Figure 6). However, a greater number of expression variants, as defined by a combination of z-score and percentile thresholds (z score −2/+2 and percentile 2.5/97.5, respectively), are observed per sample compared to other variant categories (Supplementary Figure 7; p<0.01). This highlights an opportunity for greater focus on expression variants and their clinical or biological significance.

### PORI identifies therapeutically relevant alterations in a cholangiocarcinoma patient

To demonstrate the use of PORI for clinically relevant interpretation of individual patient data, we analysed a case of cholangiocarcinoma, which was previously described as harbouring a fusion involving the oncogene *NRG1^41^*. The patient, a 38-year old woman diagnosed with intrahepatic cholangiocarcinoma, had received chemotherapy with gemcitabine and cisplatin and undergone surgery, without disease control. A metastatic tumour sample was obtained from the liver and analysed using whole-genome and transcriptome sequencing, revealing small mutations, copy number changes, structural variants, and gene expression alterations. An *ATP1B1-NRG1* gene fusion was identified which led to the rationalization for treatment with the ErbB family tyrosine kinase inhibitor afatinib, with dramatic subsequent clinical response^41^. The data from this analysis, which was processed with PORI, including matching to graphKB and display in IPR, has been made available at https://pori-demo.bcgsc.ca (IPR, PATIENT0 biop2).

PORI clearly identifies the key targetable alteration, the *NRG1* fusion, on the summary page (Supplementary Figure 8) based on matching to therapeutically relevant statements in GraphKB, along with the display of a figure describing the structural variant^42^ (Supplementary Figure 9). In addition, mutations in tumour suppressors *TP53* and *CDKN2A* are highlighted, along with amplification of the oncogenes *NTRK1* and *MCL1*. A number of genes have notably increased expression, including *NRG1*, consistent with the oncogenic effect of the gene fusion, and expression information can be viewed, sorted and filtered within IPR (Supplementary Figure 10). Details of the GraphKB associations provide information on drug sensitivity, resistance, and eligibility for clinical trials, as well as tumour type information and links to the source data in GraphKB. This provides critical support for an informed decision about therapy options, including in this case the potential for sensitivity to afatinib, which was accessed and resulted in a clinical response for this patient. In addition to specific gene associations, mutation signatures analysis^43^ (Supplementary Figure 11) reveals that this sample harbours evidence of exposure to platinum therapy (SBS31 and DBS5). This was not reported in the original published case description but is consistent with the treatment history of the patient. PORI provides flexibility for the addition of other data types when available from an analysis pipeline, including expression correlation (Supplementary Figure 12), immune environment, and mutation burden. The interactive nature of IPR allows the user to quickly view the genomic events associated with the strongest evidence of clinical relevance, and to also access the level of detail that is most pertinent, supporting informed treatment decision-making for precision oncology.

## Discussion

The rapid development of genomic technologies and bioinformatic research represents a significant challenge for precision oncology^31^. Platforms and pipelines must be able to readily incorporate new and varied content. PORI addresses this with modular reports where sections corresponding to particular specialized analyses can be added or removed as available. Previous reporting solutions have required users to input raw data and use the bioinformatic analysis pipeline integrated into the tool itself^17^. This is a barrier to use for many institutions which have already developed their own mature bioinformatic pipelines. It also limits the ability of the user to modify the pipeline as new tools are developed and new data types are added. PORI overcomes this by requiring inputs post-variant calling. Automation often comes at the cost of fine-grained control over the product. While fully automated solutions have shown promise for very common cancer types, their success with less common cancers has demonstrated there is still a strong need for human expert intervention^44^. PORI balances this by generating a fully automated report which can be manually altered and supplemented as needed.

The importance of an open-source platform is three-fold. Firstly, due to the flexible design of PORI, users will be able to contribute new content as needed both in the form of new loaders for GraphKB and new sections for new analysis types in IPR. Community involvement will help ensure the reporting platform continues to support relevant inputs that reflect the needs of the community. Secondly, the provision of a transparent option for reporting genomic data will provide an opportunity to standardize and improve reporting across multiple centres which will facilitate simpler comparisons. This is particularly critical as it has been shown that commercial platforms provide diverse results that are less amenable to scrutiny^15^. Finally, this will provide access to institutions and centres which might find the commercial alternatives cost-prohibitive.

Although PORI represents an important first step in creating an open-source standard tool for reporting in precision oncology, there are still many avenues for future development. Currently, the platform focuses on creating research reports and future iterations could include clinically accreditable report variants. Work is currently underway to create germline and pharmacogenomic report variants. Finally, perhaps the most exciting area for future work is in the application of data captured from user actions during the analysis process to iteratively improve and further automate future analysis. Facilitating the complex analysis associated with precision oncology in cancer will not only have direct benefit to the patients analyzed but also the process as a whole through improved communication and transparency.

## Methods

### GraphKB Transformation of sources for Knowledge Base Comparison

#### Import into GraphKB

Data is imported into GraphKB via automated scripts which can be found in our loader repository (Supplementary Table 4). While there is some logic specific to each source, in general the logic is that ontology terms are imported from multiple sources. Cross reference links are imported where defined and the ontology that defines the linkage is set to the source of the link. Knowledge bases are imported after ontologies as many of them require the ontologies as dependencies. For terms referenced in a knowledge base from a particular ontology the statement is linked to the specified ontology. If an ontology was not given then the term is matched by exact name match or an error is reported (Supplementary Table 3).

To ensure this process is traceable and repeatable each ontology field is stored with four main inputs: source, sourceId, name, and sourceIdVersion. The source is the ontology it was imported from (ex. HGNC). The sourceId is the Id defined by the source, this should be unique within the source (ex. 6407). The name is the human readable name of the term (ex. KRAS). and finally the sourceIdVersion is the version number of that Id. This field is optional. In some cases this may be the same as the version number of the entire resource but in many the IDs themselves are versioned independently (ex. ensembl transcript versions).

### Processing of Resources for Ontology Term Name Comparisons

The set of unique drug (or disease) names defined by each resource as well as any synonyms or product names was taken. These have been transformed to lowercase and trimmed.

#### ClinicalTrials.gov Clinical Trials

The full XML records for all trials (346,614) stored in the ClinicalTrials.gov database were downloaded on 2020-07-23 from https://clinicaltrials.gov/AllPublicXML.zip. From these, conditions and interventions were parsed into a list of terms and the frequency amongst trials of these terms. Normalization of the terms was limited to stripping trailing and leading whitespace and lowercasing. This resulted in 116,237 therapy terms and 86,204 disease terms from 345,760 clinical trials.

#### NCIt

The plain text download version of the NCIt thesaurus (Supplementary Table 2) was downloaded from NCIt (https://evs.nci.nih.gov/ftp1/NCI_Thesaurus). Terms were classified as disease or therapy based on their semantic type. Terms with the following semantic types were considered therapeutic terms (87,427 total; 5,017 primary): Antibiotic; Biologically Active Substance; Biomedical or Dental Material; Chemical Viewed Functionally; Chemical Viewed Structurally; Chemical; Clinical Drug; Drug Delivery Device; Element, Ion, or Isotope; Food; 'Hazardous or Poisonous Substance; Hormone; Immunologic Factor; Indicator, Reagent, or Diagnostic Aid; Inorganic Chemical; Medical Device; Organic Chemical; Pharmacologic Substance; Plant; Steroid; Substance; Therapeutic or Preventive Procedure; and, Vitamin. Terms with the following semantic types were considered disease terms (57,276 total; 6,526 primary): Anatomical Abnormality; Congenital Abnormality; Disease or Syndrome; Experimental Model of Disease; Mental or Behavioral Dysfunction; Neoplastic Process; Sign or Symptom. Both the names and synonyms of the terms were considered.

#### DrugBank

DrugBank ^29^ was downloaded in its XML format (for version information, see Supplementary Table 2) from https://go.drugbank.com/releases. Names were extracted from the records based on the name, synonyms, and products tags. This resulted in 131,412 unique terms.

#### FDA SRS

The UNII identifiers (Supplementary Table 2) were downloaded from the FDA substance registration system (FDA SRS) at https://fdasis.nlm.nih.gov/srs/download/srs/UNII_Data.zip. The PT field was used as the name field. This resulted in 109,334 primary terms (all terms have unique identifiers and therefore are considered non-alias primary terms).

#### Disease Ontology

The disease ontology^28^ was downloaded as a JSON (Supplementary Table 2) from https://github.com/DiseaseOntology/HumanDiseaseOntology. Terms were extracted from the lbl and synonyms attributes. This resulted in 19,064 primary terms out of 38,489 total terms.

#### ChEMBL

The postgres dump of the ChEMBL^30^ database was downloaded (https://chembl.gitbook.io/chembl-interface-documentation/downloads) and a plain text version of the drug names was created from the molecule_dictionary and molecule_synonym tables. Where the preferred name field of the first table was used as the primary set of terms and names from the synonyms table were included in the full set (Supplementary Table 2). This resulted in 35,219 primary terms out of 123,287 total terms.

### Processing of cBioportal.org TCGA Data

All TCGA pan-cancer ATLAS data was downloaded from cBioportal.org^39^. This consisted of the all *_tcga_pan_can_atlas_2018 studies: blca, brca, cesc, chol, coadread, dlbc, esca, gbm, hnsc, kich, kirc, kirp, laml, lgg, lihc, luad, lusc, meso, ov, paad, pcpg, prad, sarc, skcm, tgct, thca, thym, ucec, ucs, and uvm.

Variants were compiled for each sample (includes some repeat samples from the same patient) from the expression (data_RNA_Seq_v2_mRNA_median_all_sample_Zscores.txt); copy number (data_CNA.txt); small mutations (data_mutations_extended.txt); and fusions (data_fusions.txt) files. Copy variants were classified as deep deletion with a value of −2 or lower and amplification with a value of 2 or greater. The distribution of copy number values amongst all patients was plotted as a sanity check that these values would result in outliers and not represent a large percentage of the calls (Supplementary Figure 13). The expression z-scores are pre-calculated in the cBioportal data. A threshold combination of z-score (−2, +2) and percentile (2.5, 97.5) was used in evaluating expression variants to determine outliers. Small mutations and fusion matching was limited to genes and mutations with protein changes (for small mutations). For simplicity, matching did not include intergenic mutations.

The number of variants called per patient was compared across all studies for each variant type (Supplementary Figure 7) which had median values of: 402 expression variants (exp); 1 gene fusion (fus); 84 copy variants (cnv); and 53 small mutations (mut). A one-way ANOVA was performed to determine significance (F-statistic: 4613.24, p-value: 0.0) followed by a TukeyHSD test to investigate pairs. The null hypothesis was rejected for all combination pairs (FWER=0.05, adjusted p-value of 0.001) with the exception of fusions and small mutations which had a p-value of 0.9.

Each variant was then matched to GraphKB using the GraphKB python adaptor (Supplementary Table 4). From these statement matches, the number of matches by variant type per each conclusion was determined. The conclusion of a statement is considered to be the combination of its relevance and subject fields (Figure 4).

## Code Availability

The implementation and peer review details of code for PORI can be found in https://github.com/bcgsc/pori and related repositories (Supplementary Table 4).

## Data Availability

The data that support the findings of this study are available from multiple third parties but restrictions apply to the availability of some of these data, which were used under license for the current study. Data are however available from the authors upon reasonable request and with permission of the applicable third party

## Acknowledgements

This work would not be possible without the participation of our patients and families, the POG team, the GSC platform, and the generous support of the BC Cancer Foundation and Genome British Columbia (project B20POG). We also acknowledge contributions towards equipment and infrastructure from Genome Canada and Genome BC (projects 202SEQ, 212SEQ, 12002), Canada Foundation for Innovation (projects 20070, 30981, 30198, 33408 and 35444), the BC Knowledge Development Fund, and the Canada Research Chairs program to SJMJ. The results published here are in part based upon data generated by the following projects and obtained from dbGaP (http://www.ncbi.nlm.nih.gov/gap): The Cancer Genome Atlas managed by the NCI and NHGRI (http://cancergenome.nih.gov); Genotype-Tissue Expression (GTEx) Project, supported by the Common Fund of the Office of the Director of the National Institutes of Health (https://commonfund.nih.gov/GTEx).

