## Supplement for "A Platform for Oncogenomic Reporting and Interpretation"

### Supplementary Data

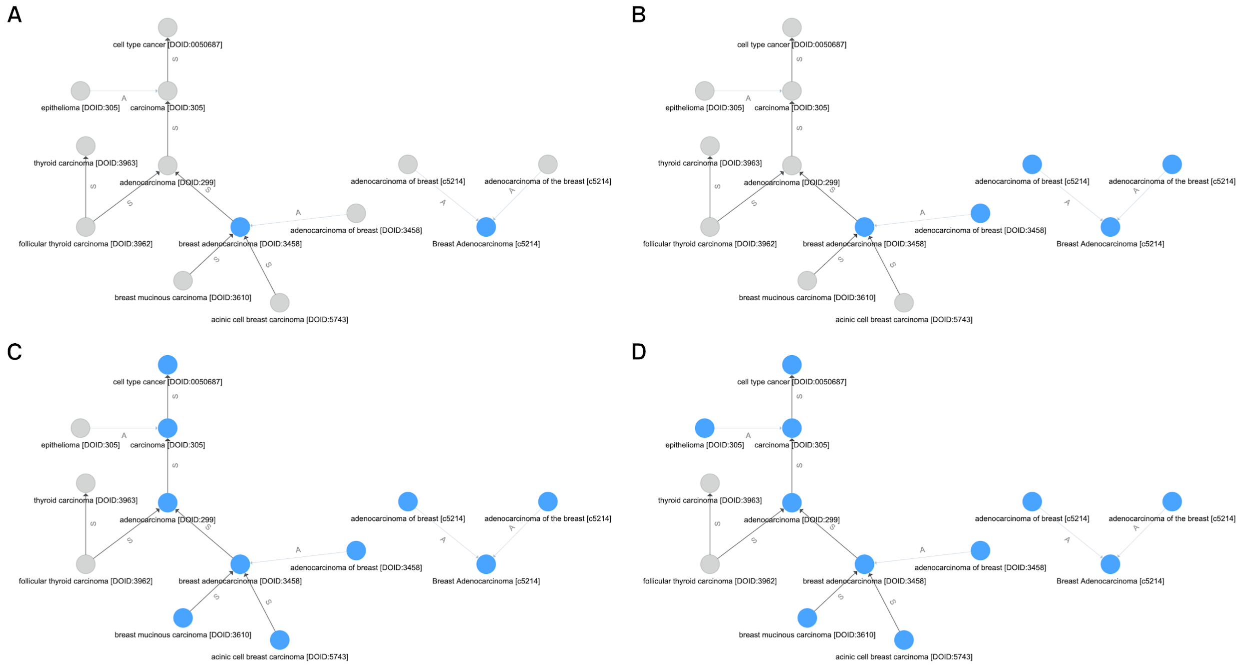


*Supplementary Figure 1. Resolving equivalent disease terms through the matching algorithm implemented in the graphkb python adaptor which leverages the graph model of GraphKB. A subset of the related terms is shown for simplicity, the input is “breast adenocarcinoma”. Matched terms are highlighted in blue. Edges labelled above are ‘subclass of’ (S) or ‘alias of’ (A) relationships. There are four steps involved in the matching algorithm: exact matching by name (A); expansion of alias and deprecated terms (B); gathering ancestors and descendants along the subclass tree (C); and finally re-expansion of aliases and deprecated terms from newly collected parent and child terms (D).*

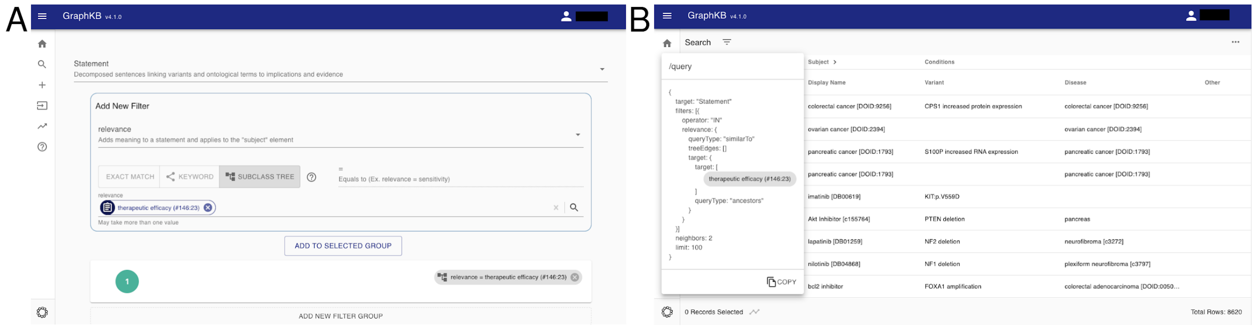


*Supplementary Figure 2. Queries (A) built in the web client display the corresponding API call (B) to aid users in familiarizing themselves with querying via the API.*

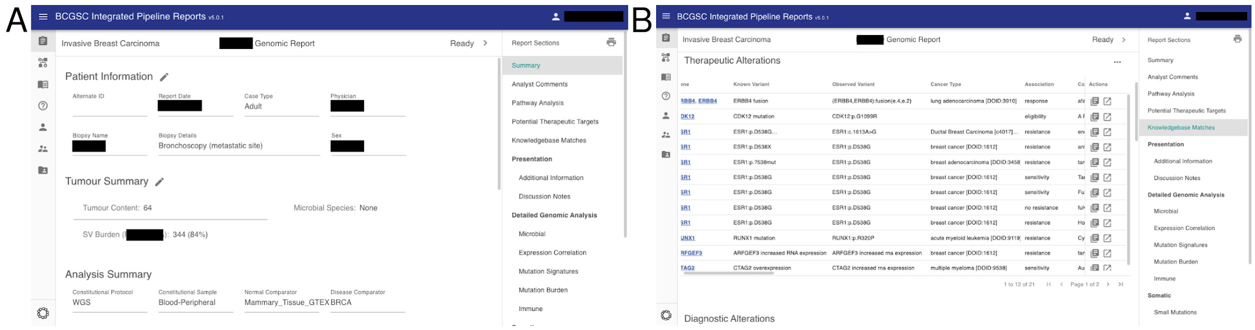


*Supplementary Figure 3. The report front page (A) view in the IPR web application. The annotations collected from GraphKB are listed in the knowledge base matches section (B) section of the generated report with links from each match back to its corresponding statement in GraphKB.*

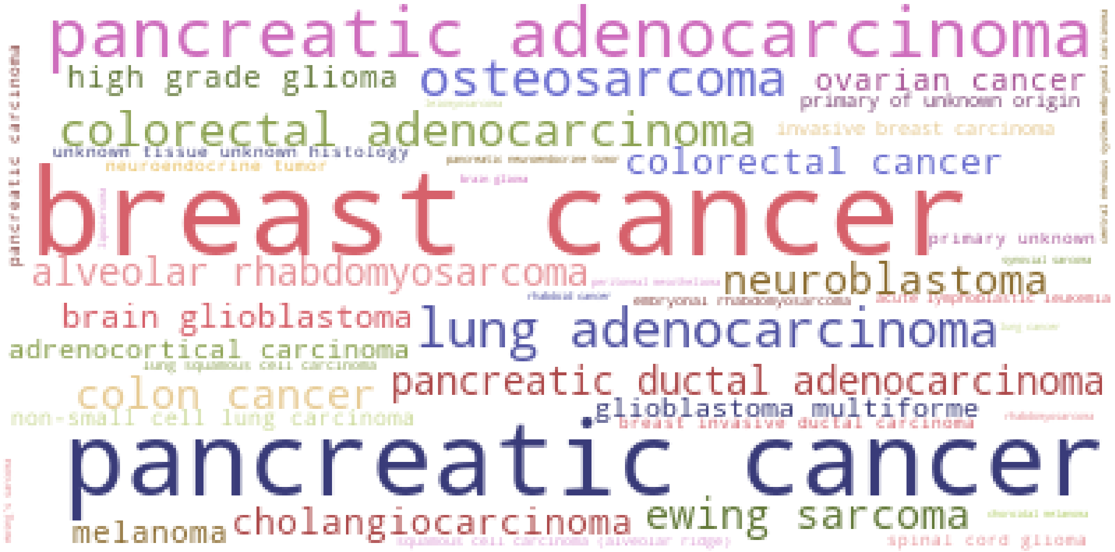


*Supplementary Figure 4. Diagnosis distribution of the 440 patients with 171 different diagnoses analyzed using PORI through the POG program.*

*Supplementary Table 1. Ontology or Resource Choices for Controlled Vocabulary in Common Cancer Knowledge bases*

| KB Name | Genes | Drugs | Diseases |
|---|---|---|---|
| CIViC | Entrez Gene | NCIt | Disease Ontology |
| OncoKB | Entrez Gene | NCIt | not specified |
| CGI | not specified | not specified | not specified |
| COSMIC* | custom | not specified | not specified |
| MetaKB | HGNC | ChEMBL | Disease Ontology |
| JAX-CKB | HGNC | not specified | Disease Ontology |
| PMKB | HGNC | N/A | custom |
| My Cancer Genome | RefSeq | NCIt | NCIt |
| CanDL | HGNC | N/A | not specified |

* Download of the resistance mutations data

*Supplementary Table 2. Disease and Drug Definition Resources*

| Resource | Resource Version | Primary Terms | Total Terms (+) |
|---|---|---|---|
| Disease Ontology | v2020-06-18 | 19064 | 38489 |
| NCIt (diseases) | v20.06e | 6526 | 57276 |
| OncoTree | 2020_04_01 | 851 | 851 |
| **Total Diseases** | | **25063** | **87687** |
| DrugBank | v5.1.7 | 13599 | 129160 |
| FDA SRS | 27Mar2020 | 109334 | 109334 |
| NCIt (drugs) | v20.06e | 5017 | 87427 |
| ChEMBL | chembl_27 | 35219 | 123287 |
| **Total Drugs** | | **137216** | **391828** |

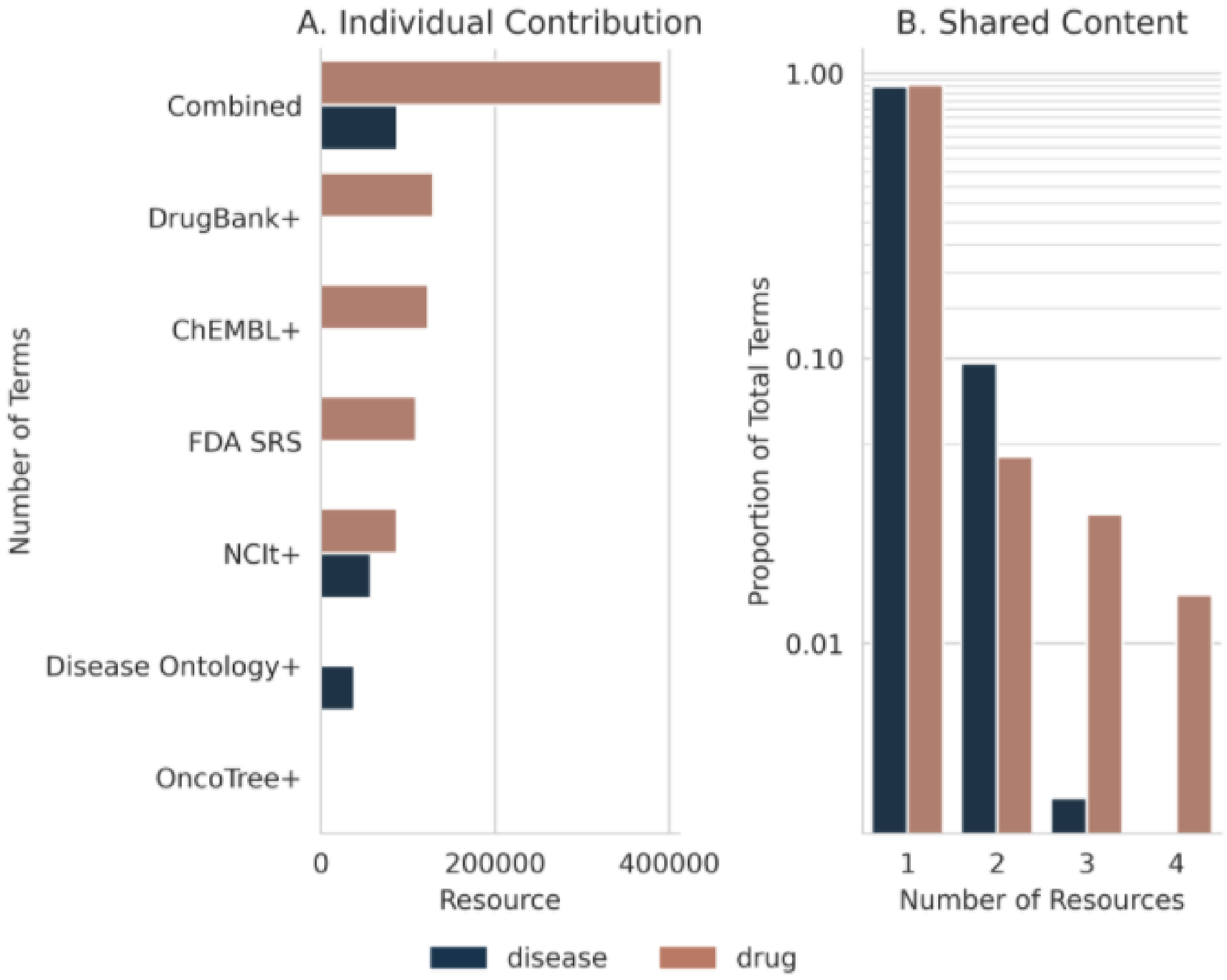


*Supplementary Figure 5. Number of terms shared between resources for disease and drug terms. Most terms are unique to a given resource (diseases: 0.90, drugs: 0.91). The '+' is used to indicate that all terms, including deprecated forms and aliases, were included in this analysis. The relative sizes of the individual resources is given (A) as well as the proportion of terms which were common amongst a minimum number of resources (B).*

*Supplementary Table 3. Loading Percentages of knowledge base Sources. Records is the number of records processed without error by the number of total records in the original source format. The statement rate success is a measure of the number of statements these records could generate by the number of statements that were successfully created.*

| Source | Date Accessed | Records | Statements | % Success |
|---|---|---|---|---|
| CGI | 2019-05-24* | 1647 | 966 / 1647 | 58.7 |
| CIViC | 2020-10-12 | 3260 | 3346 / 3579 | 93.4 |
| COSMIC | 2020-10-18 | 4100 | 1703 / 1703 | 100 |
| DoCM | 2020-10-19 | 1364 | 8615 / 8627 | 99.9 |
| OncoKB | 2020-07-05 | 5165 | 9126 / 9234 | 98.8 |

* This resource was not re-downloaded at a later date as the data has not been updated, last checked 2020-02-17

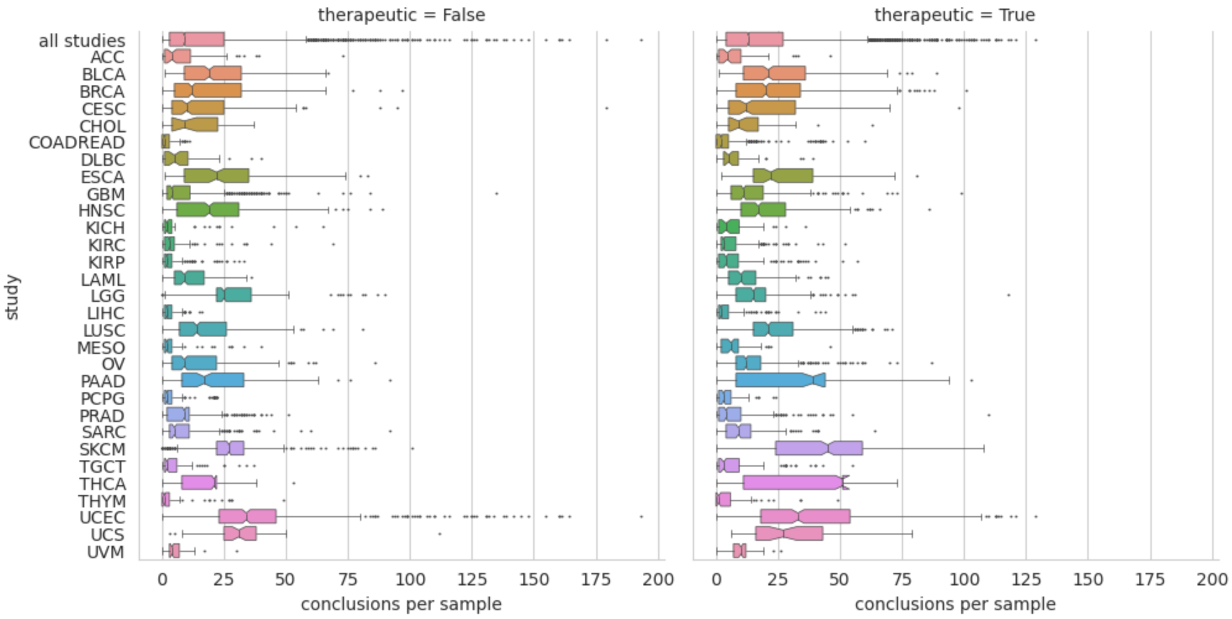


*Supplementary Figure 6. Number of therapeutic (right) and non-therapeutic (left) unique conclusions per sample for TCGA samples downloaded from cbioportal.org. The combined set of all studies is shown as the top bar “all studies”. Box plots represent the median, upper and lower quartiles of the distribution, and whiskers represent the limits of the distribution (1.5-times interquartile range).*

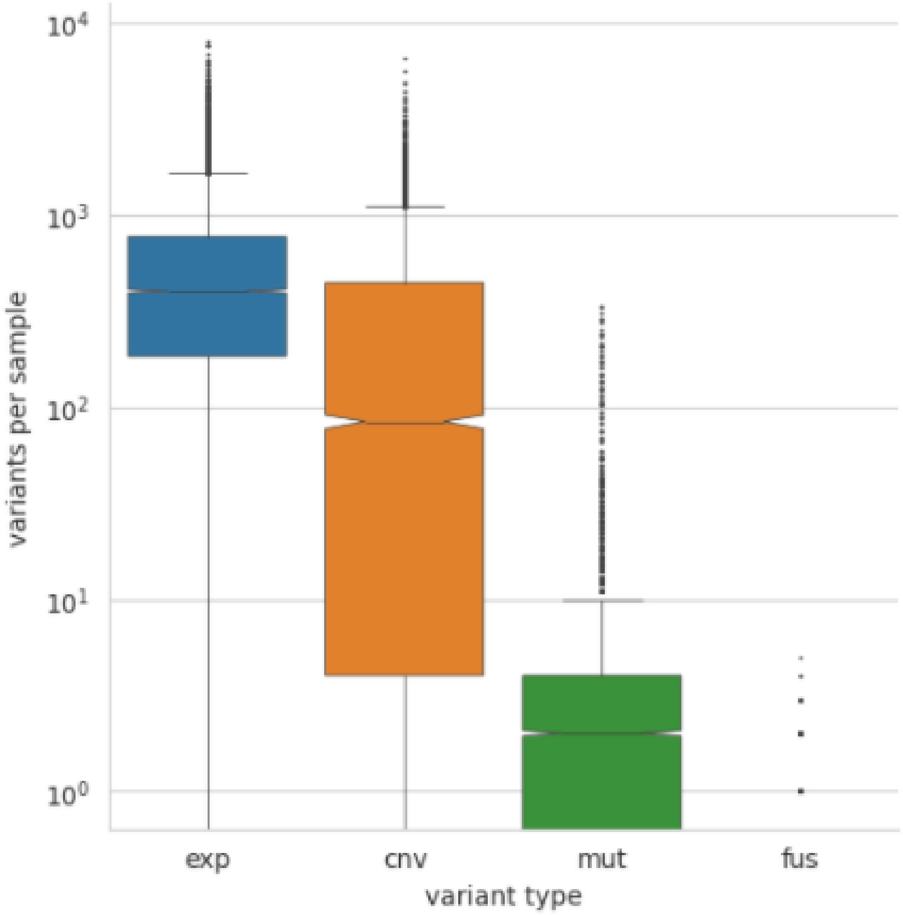


*Supplementary Figure 7. Number of variant calls per sample. The median number of variants called per sample for each variant type is: 402 expression variants (exp); 0 gene fusions (fus); 84 copy variants (cnv); and 2 small mutations (mut). Box plots represent the median, upper and lower quartiles of the distribution, and whiskers represent the limits of the distribution (1.5-times interquartile range).*

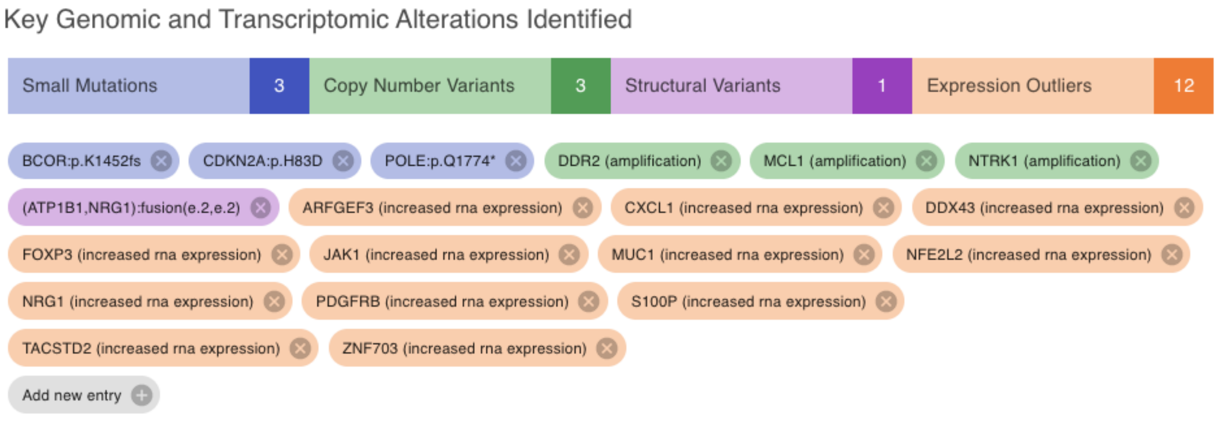


*Supplementary Figure 8. Key alterations automatically identified on the PORI report for POG case (Demo PATIENT0 biop2). The actionable fusion is shown in purple.*

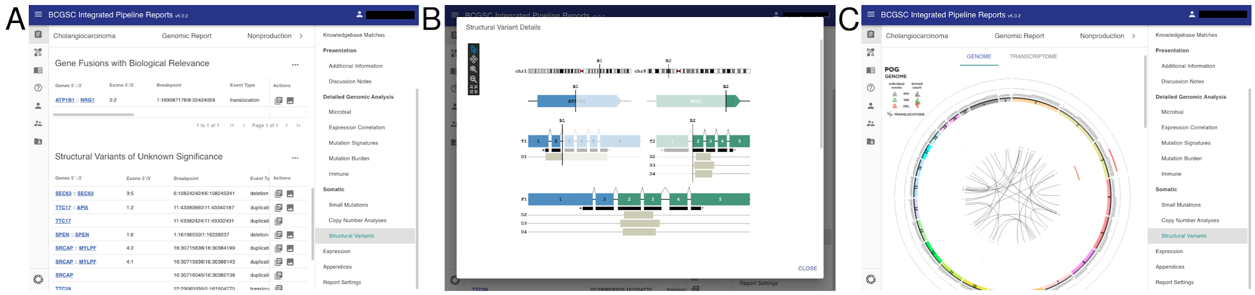


*Supplementary Figure 9. The structural variants section of IPR provides additional details on structural variants reported in the current patient. Details can be explored in the interactive tables (A). MAVIS fusion diagrams (or user's choice) can be shown in-line with the details of the structural variant using the image button in the action columns (B). Circos plots may be loaded with the report as well to show a comprehensive quick overview of the total landscape of structural variants (C).*

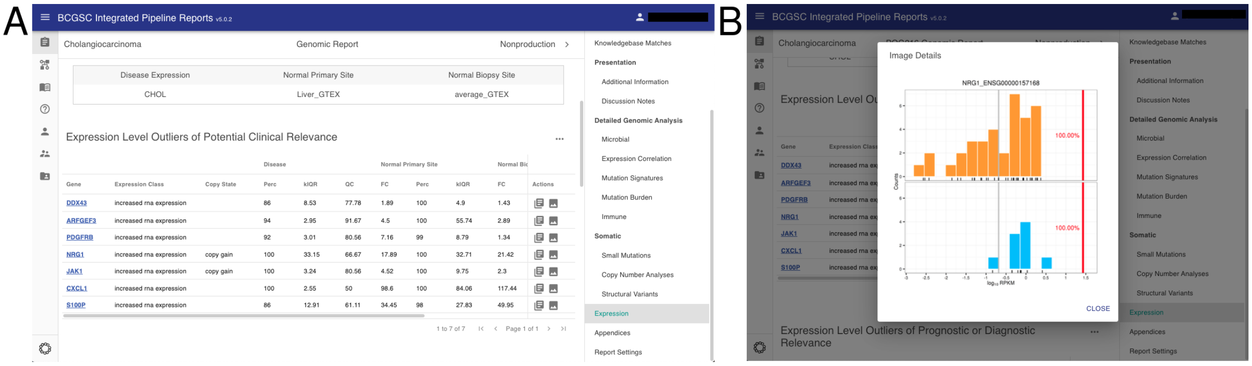


*Supplementary Figure 10. Expression data is displayed in tables which can be sorted and filtered (A). Additionally images showing further details regarding the distribution of this patient relative to the expression of other samples within the selected comparator cohort can be displayed using the image button in the actions column of the table (B).*

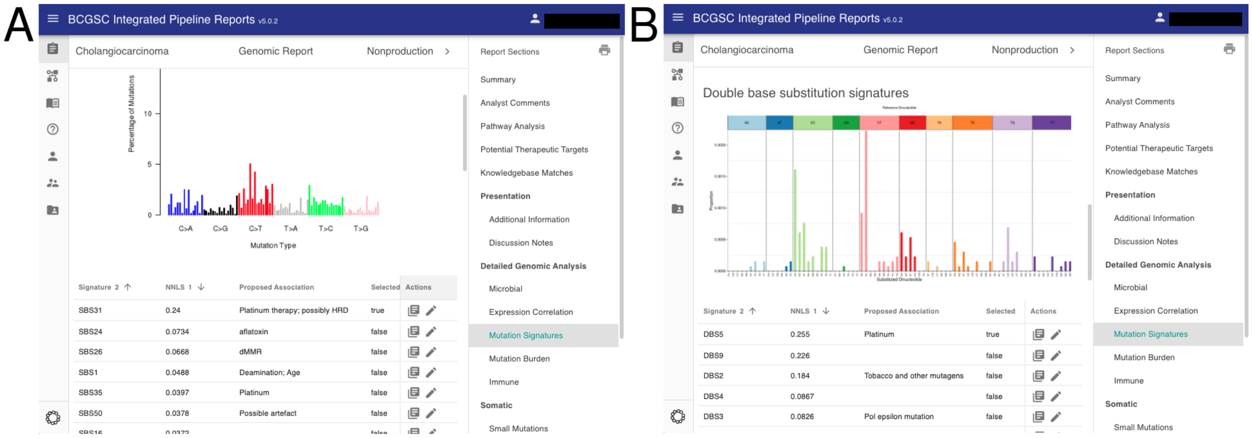


*Supplementary Figure 11. The mutation signatures section reports non-negative least squares values for mutation signatures computed for the current patient. These include single base substitution signatures (A); double base substitution signatures (B); and indel signatures.*

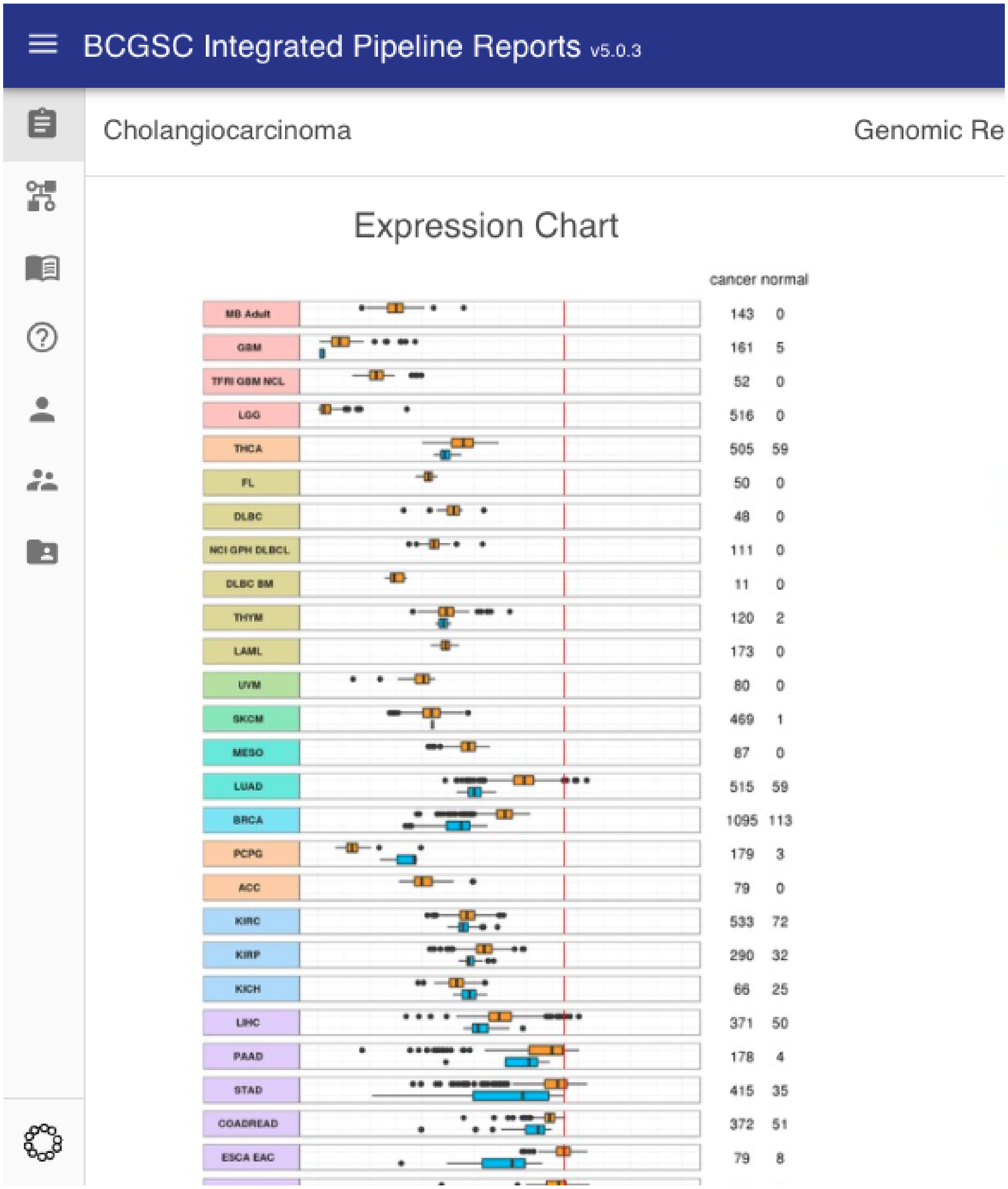


*Supplementary Figure 12. Spearman Correlation of RNA expression values of the current patient against expression values from TCGA and other disease specific datasets. The vertical red line indicates this patient's value. For each cohort, values are split by disease status (cancer versus normal). This plot is expected to show correlation with the primary site of the given tumour type, here cholangiocarcinoma (CHOL).*

*Supplementary Table 4. Links to all open source content. Versions listed are those used for this manuscript*

| **Component** | **Version** | **Repository** |
|---|---|---|
| Full Platform | | https://github.com/bcgsc/pori |
| IPR adapter | 2.0.4 | https://github.com/bcgsc/pori_ipr_python |
| GraphKB adapter | 1.5.1 | https://github.com/bcgsc/pori_graphkb_python |
| GraphKB Loader | 5.0.0 | https://github.com/bcgsc/pori_graphkb_loader |
| GraphKB client | 4.2.1 | https://github.com/bcgsc/pori_graphkb_client |
| GraphKB API | 3.13.0 | https://github.com/bcgsc/pori_graphkb_api |
| IPR client | 6.0.4 | https://github.com/bcgsc/pori_ipr_client |
| IPR API | 6.0.0 | https://github.com/bcgsc/pori_ipr_api |
| GraphKB schema | 3.14.3 | https://github.com/bcgsc/pori_graphkb_schema |
| GraphKB parser | 1.1.1 | https://github.com/bcgsc/pori_graphkb_parser |
| PORI cBioportal | | https://github.com/bcgsc/pori_cbioportal |

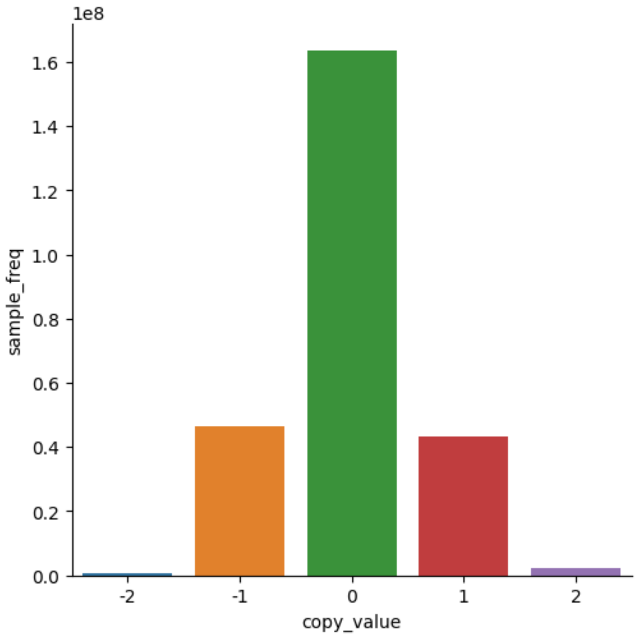


*Supplementary Figure 13. Distribution of copy number values amongst all patients in the TCGA 2018 discrete copy number data downloaded from cbioportal.org. Values of -2 were called as deep (homozygous) deletions and values of +2 were called as amplifications.*
